# Anatomical characterisation of somatostatin-expressing neurons belonging to the anterolateral system

**DOI:** 10.1101/2024.11.19.624289

**Authors:** Wenhui Ma, Erika Polgár, Allen C. Dickie, Mai Abu Hajer, Raphaëlle Quillet, Maria Gutierrez-Mecinas, Mansi Yadav, Junichi Hachisuka, Andrew J Todd, Andrew M. Bell

**Affiliations:** School of Psychology and Neuroscience, Sir James Black Building, University of Glasgow, Glasgow, G12 8QQ, UK; School of Biodiversity, One Health and Veterinary Medicine, University of Glasgow, G61 1QH, UK

## Abstract

Anterolateral system (ALS) spinal projection neurons are essential for pain perception. However, these cells are heterogeneous, and there has been extensive debate about the roles of ALS populations in the different pain dimensions. We recently performed single-nucleus RNA sequencing on a developmentally-defined subset of ALS neurons, and identified 5 transcriptomic populations. One of these, ALS4, consists of cells that express *Sst*, the gene coding for somatostatin, and we reported that these were located in the lateral part of lamina V. Here we use a Sst^Cre^ mouse line to characterise these cells and define their axonal projections. We find that their axons ascend mainly on the ipsilateral side, giving off collaterals throughout their course in the spinal cord. They target various brainstem nuclei, including the parabrachial internal lateral nucleus, and the posterior triangular and medial dorsal thalamic nuclei. We also show that in the L4 segment *Sst* is expressed by ∼75% of ALS neurons in lateral lamina V and that there are around 120 *Sst*-positive lateral lamina V cells on each side. Our findings indicate that this is a relatively large population, and based on projection targets we conclude that they are likely to contribute to the affective-motivational dimension of pain.

## INTRODUCTION

The spinal dorsal horn receives somatosensory information from primary afferents that innervate the skin and deep tissues, and encode various stimulus modalities. This information is processed through complex neural circuits before being transmitted to the brain through spinal projection neurons^1–4^. Many of these belong to the anterolateral system (ALS), which consists of cells whose axons innervate a distinctive set of brain regions, including thalamus, periaqueductal grey matter (PAG), superior colliculus, lateral parabrachial area (LPB), nucleus of the solitary tract (NTS) and brainstem reticular formation^5–11^. Cell bodies of ALS neurons are found in lamina I, the deeper laminae (III-VIII), the area around the central canal (lamina X), and the lateral white matter^12–16^. The majority of ALS neurons project to the brain on the contralateral side, although there are also neurons that project bilaterally or only to the ipsilateral side. In humans, ascending ALS axons are located in the anterolateral funiculus, and disruption of these by anterolateral cordotomy results in loss of pain, itch and skin temperature sensation on the opposite side of the body caudal to the lesion^17,18^, demonstrating the importance of the ALS for these sensory perceptions. In rodents, ALS axons are located more dorsally, within the lateral funiculus^10,11,19–21^, presumably because the corticospinal tract (which occupies the dorsal part of the lateral funiculus in humans) is located in the dorsal columns.

ALS neurons are functionally heterogeneous, and there has been extensive debate about the relative importance of different populations, for example those that are located in deep or superficial laminae, in transmitting pain information^12,22–26^. However, the lack of molecular markers for different subpopulations of ALS cells has made it difficult to target specific types of ALS neuron selectively, and this has limited our ability to define the roles of the different cell types. We recently performed single-nucleus RNA sequencing (snRNAseq) on a developmentally-defined subset of ALS cells (those that transiently express the transcription factor Phox2a^9^), and identified 5 populations, which we named ALS1-5^27^. One of these populations, ALS4, consisted of cells that were mainly located in the lateral reticulated part of lamina V^28,29^. Many of the ALS4 cells contain *Sst* mRNA, which codes for the peptide somatostatin, and this was relatively specific for this population. Here we have used a mouse line in which Cre recombinase is expressed from the *Sst* locus to selectively target and characterise the ALS4 cells. We tested the hypothesis that these transcriptomically-defined cells have characteristic patterns of axonal projections that may dictate their distinct roles in the generation of pain.

## RESULTS

### Anterograde tracing in the Sst^Cre^ mouse

To reveal axonal projections of *Sst*-expressing ALS neurons we injected AAVs coding for Cre-dependent tdTomato into either the L3 or L3, L4 and L5 spinal segments of Sst^Cre^ mice (Table 1). Due to the very high density of *Sst*-expressing dorsal horn neurons^30–32^ there was extensive tdTomato labelling close to the injection site; however, at more rostral levels of the spinal cord labelling was much more restricted (Fig 1). At upper lumbar levels, there was a prominent bundle of fibres within the ipsilateral ventral funiculus, and axons could be seen entering this region near the injection site. In thoracic segments, this bundle lay within the lateral funiculus and it ascended through the entire length of the spinal cord, moving more dorsally as it did so, as indicated by the arrows in Fig 1. A much smaller bundle (double arrows in Fig 1) was present in the same position on the contralateral side. Labelled fibres and varicosities were also present in the lateral spinal nucleus (LSN) on the ipsilateral side in lumbar and thoracic segments (arrowheads in Fig 1), but the density of LSN labelling decreased progressively, and it was virtually absent in cervical segments. These LSN axons are likely to originate from *Sst*-expressing excitatory interneurons in the superficial dorsal horn (SDH), as we have seen a similar projection to the LSN when AAV.Cre^ON^.GFP was injected into the lumbar spinal cord of Tac1^Cre^ mice, and many of the varicosities in the LSN seen in that study contained somatostatin^21^, consistent with extensive coexpression of these peptides in SDH excitatory interneurons^31,33^. At all spinal levels examined, collateral branches with boutons occupied a region extending from the lateral reticulated part of lamina V towards the central canal, and these appeared to originate from the main fibre bundle that ascended in the lateral funiculus (Fig 1). These collaterals were particularly well seen in animals that had received injections of AAV.Cre^ON^.tdTomato into the L3, L4 and L5 segments (Fig S1). Only very sparse collaterals were seen in lamina I in segments rostral to T10.

**Fig 1.**
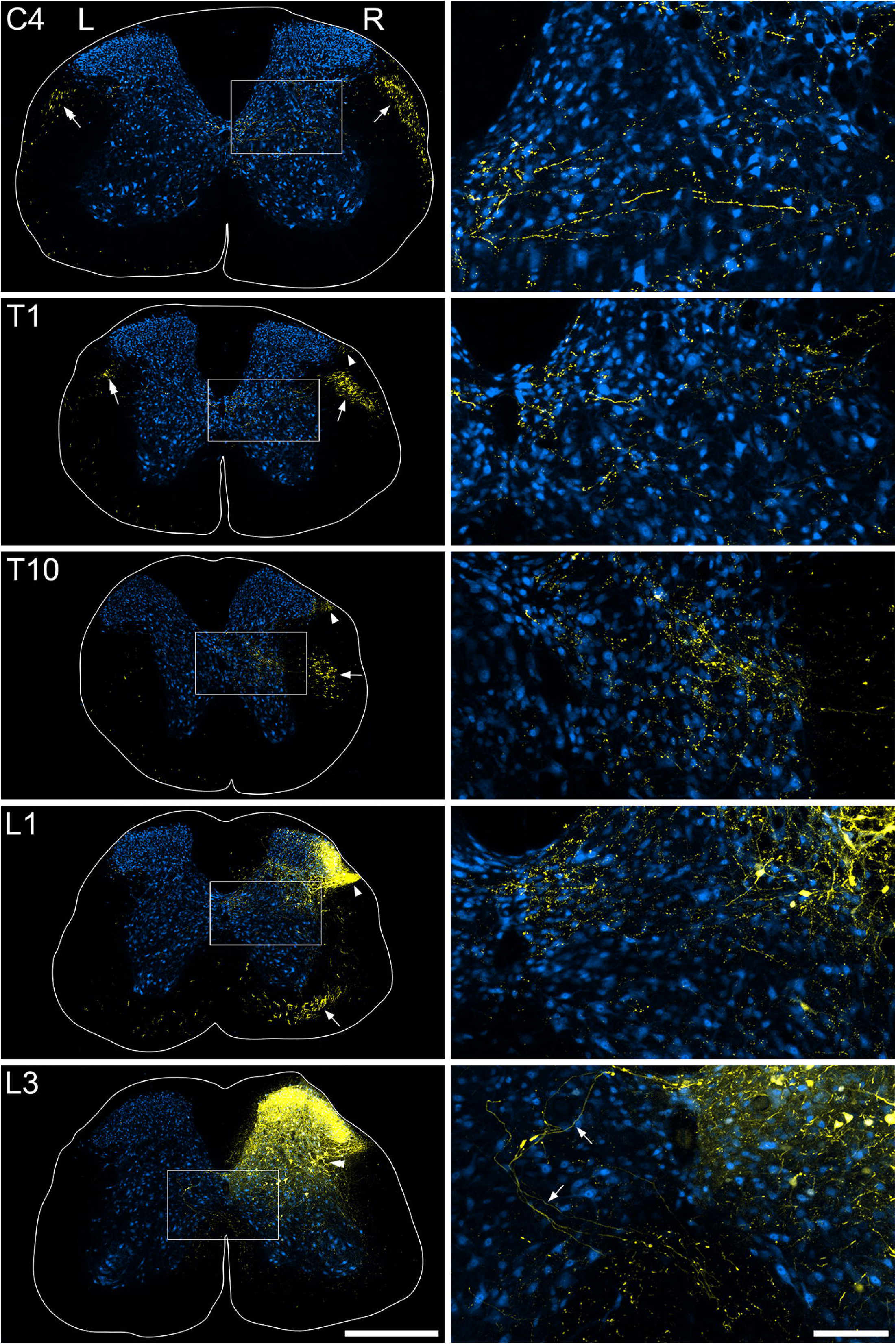
TdTomato labelling (yellow) in the spinal cord of a Sst^Cre^ mouse that had received intraspinal injection of AAV.Cre^ON^.tdTomato into the L3 segment on the right side 5 weeks prior to perfusion fixation (#4 in Table 1). Neuronal labelling with NeuN antibody is shown in blue. Images on the left are confocal scans through transverse sections at different segmental levels (C4, T1, T10, L1 and L3). Higher magnification views of the regions marked by boxes are shown on the right. At the level of the injection site (L3) there is dense labelling in the dorsal horn, including the lateral reticulated part of lamina V (double arrowheads). In the high magnification image from this section, some labelled axons can be seen passing towards the ventral funiculus on the same side (arrows). At more rostral levels, a prominent bundle of labelled axons can be seen in the ventral funiculus (L1) or lateral funiculus (T10, T1, C4), and this moves progressively more dorsally as it ascends. A much smaller bundle is visible in the same location on the opposite (contralateral) side in the C4 and T1 segments (double arrows). Many labelled structures are visible in the ipsilateral LSN in the L1 segment (arrowheads), and this labelling extends rostrally to at least the T1 segment, although it becomes less intense in rostral thoracic segments. Scale bars for main images = 500 μm, for high magnification images = 100 μm.

**Table 1.** Experiment details for Sst^Cre^ mice used in viral anterograde tracing experiments. Mice received injections into either the L3 segment only, or into the L3, L4 and L5 segments. **L**, left side; **R**, right side. In all cases coronal sections through the brain as far as the rostral part of the thalamus were examined. Transverse sections through selected spinal segments were examined, and these are listed in the 5^th^ column. For details of viruses see Table S1.

| Animal | Sex | Injection details | Survival (days) | Spinal segments examined |
| --- | --- | --- | --- | --- |
| 1 | M | AAV.Cre<sup>ON</sup>.tdTom **L** L3 | 29 | C7, T7, L1, L3 |
| 2 | M | AAV.Cre<sup>ON</sup>.tdTom **L** L3 | 28 | C7, T7, L1, L3 |
| 3 | M | AAV.Cre<sup>ON</sup>.tdTom **R** L3, L4, L5 | 35 | C1, C7, L3, L4, L5, |
| 4 | F | AAV.Cre<sup>ON</sup>.tdTom **R** L3 | 35 | C1, C4, C7, T1, T4, T7, T10, L1, L3 |
| 5 | F | AAV.Cre<sup>ON</sup>.tdTom **R** L3, L4, L5 | 54 | C4, C7, T7, L4 |
| 6 | F | AAV.Cre<sup>ON</sup>.tdTom **R** L3, L4, L5 | 54 | C7, T7, T10, T12, L4 |
| 7 | F | AAV.Cre<sup>ON</sup>.tdTom **R** L3, L4, L5 | 53 | C7, T7, T10, T12, L4 |
| 8 | F | AAV.Cre<sup>ON</sup>.tdTom **R** L3 | 53 | C7, T4, T7, T10, T12, L1, L3 |

Within the brainstem the main ascending fibre bundle was located near the ventrolateral surface on the ipsilateral side, with a much smaller bundle in the same location on the contralateral side (Fig 2). These bundles continued in the same ventrolateral location up to the rostral part of the pons (at a level corresponding to ∼interaural −1 mm in the Franklin and Paxinos atlas^34^). Here, the fibres passed dorsally and then caudally to reach the lateral parabrachial area, where they arborised mainly in the internal lateral nucleus (PBil), with much smaller contributions to other regions in LPB (Fig 3). Many fibres crossed the midline by looping caudally in the superior medullary velum (SMV) to reach the PBil of the opposite side (Fig 2). The density of labelling in PBil was nearly symmetrical between the two sides, presumably because some fibres that ascended on the ipsilateral side of the caudal brainstem reached the contralateral PBil through the SMV. In their course within the caudal brainstem, collateral branches were given off to the caudal part of the NTS, the parvicellular reticular nucleus and the gigantocellular reticular nucleus. A relatively dense bundle was given off from the main ascending tract at the level of the facial nerve and this arborised in the intermediate reticular nucleus and adjacent structures (Fig S2).

**Fig 2.**
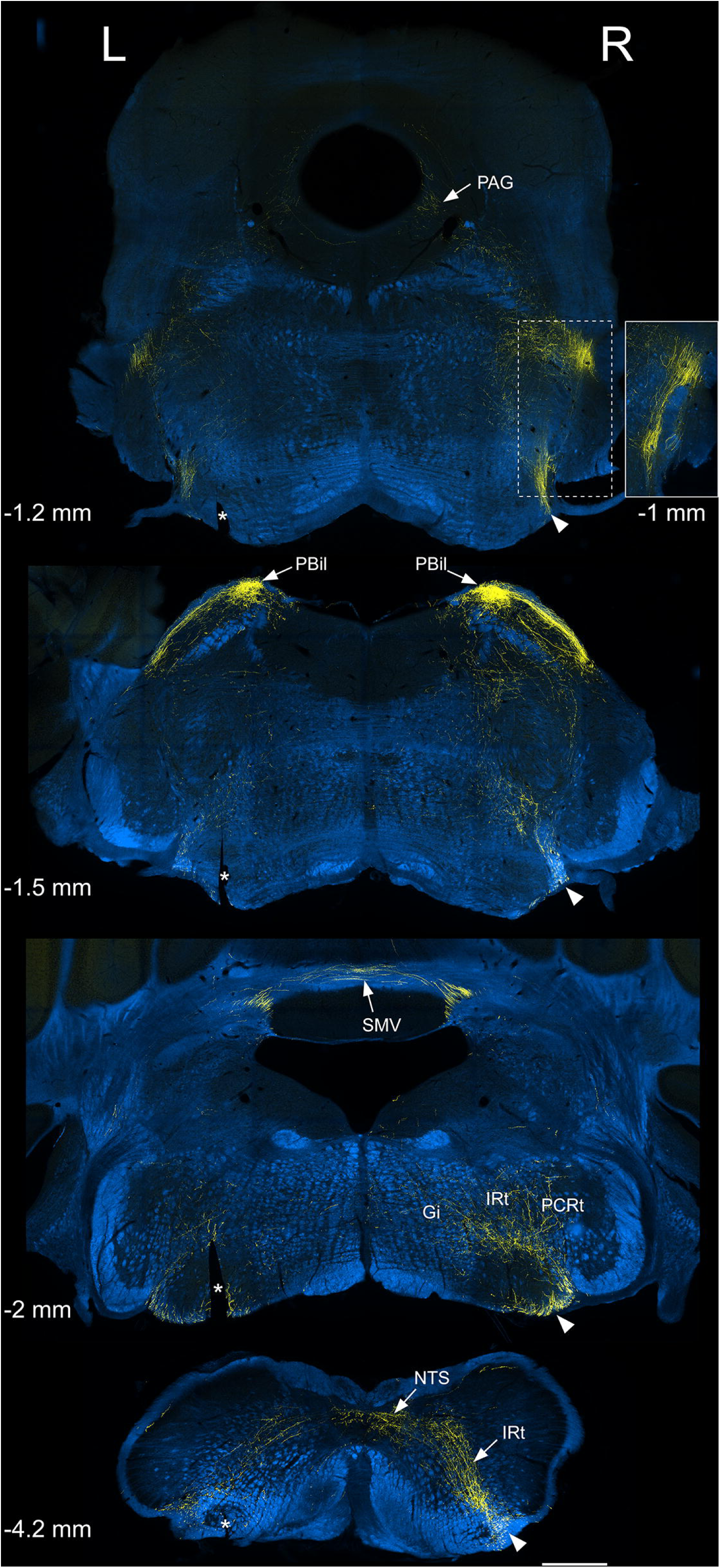
TdTomato labelling in the brainstem of the Sst^Cre^ mouse illustrated in Fig 1 (#4 in Table 1). TdTomato is shown in yellow, superimposed on images acquired with dark-field illumination (blue). Four coronal sections of the brainstem are shown, and the approximate locations in relation to the interaural line are indicated^34^. The main ascending bundle of axons is located on the ventrolateral aspect of the brainstem on the side ipsilateral to the spinal injection (Right, R), and is marked with arrowheads. Collateral branches are given off at several levels in the caudal brainstem, in particular to the NTS, Gi, IRt and PCRt. There is a dense projection to the lateral parabrachial area (LPB), which is largely restricted to the PBil. Fibres enter the LPB by passing along the dorsolateral aspect of the pons (as seen at the −1.5 interaural level), and these connect with the main ascending bundle on the ventrolateral aspect at a slightly more rostral level (as shown in the inset). A few fibres can be seen in the PAG. Note that although labelling is predominantly ipsilateral to the injection site in the caudal brainstem, there is a substantial projection to the PBil on both sides, and some of this is likely to be through fibres that cross the midline in the superior medullary velum (SMV). Asterisks indicate a cut made on the left side that was used to orientate the sections. The inset corresponds to the area marked with a dashed line on the top image and shows part of a section approximately 0.2 mm rostral (i.e. interaural −1 mm). Scale bar = 500 μm. Gi, gigantocellular reticular nucleus; IRt, intermediate reticular nucleus; NTS, nucleus tractus solitarius; PAG, periaqueductal grey matter; PBil, lateral parabrachial nucleus, internal part; PCRt, parvicellular reticular nucleus; SMV, superior medullary velum.

**Fig 3.**
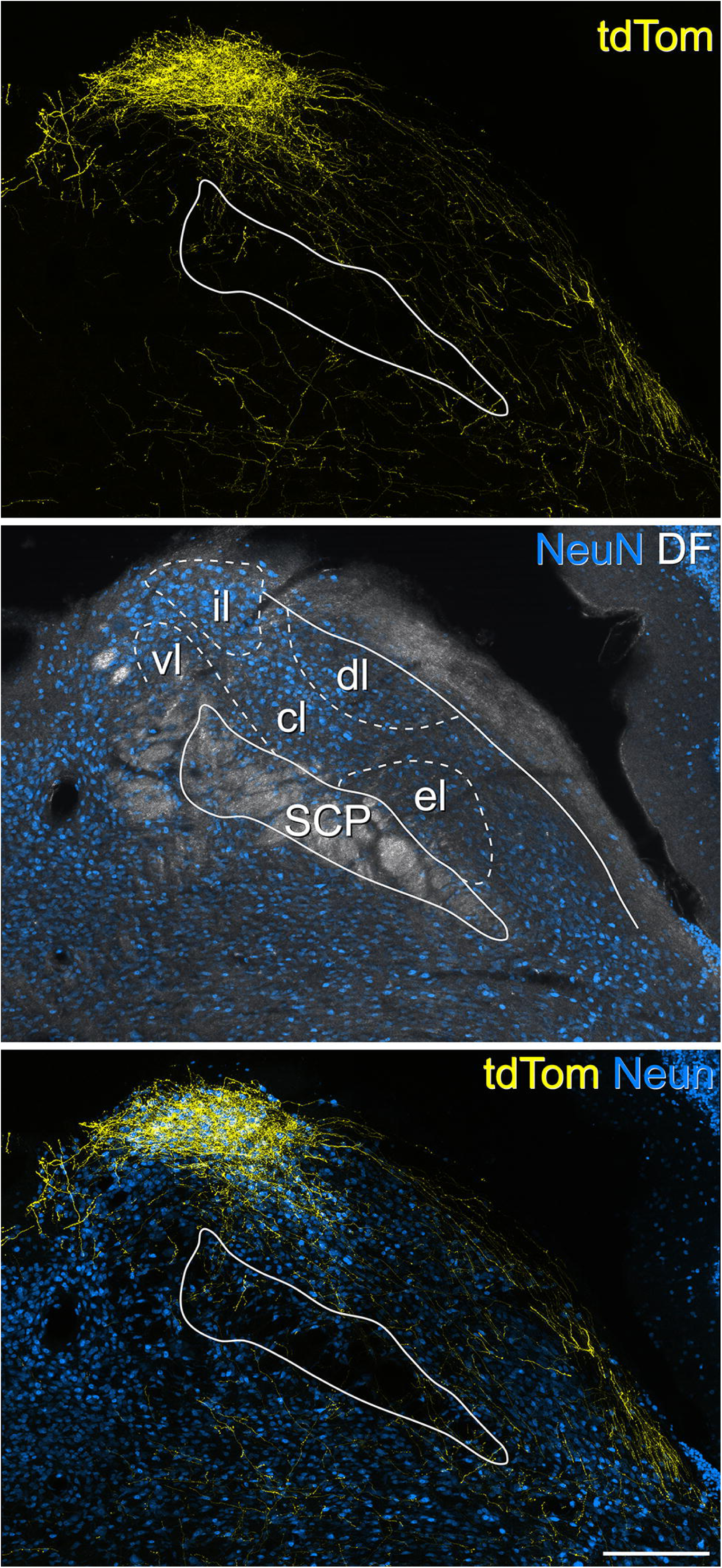
TdTomato labelling in the right lateral parabrachial area of a Sst^Cre^ mouse that had received intraspinal injection of AAV.Cre^ON^.tdTomato into the L3 segment on the right side ∼8 weeks previously (#8 in Table 1). TdTomato labelling is shown in yellow, NeuN in blue and in the middle image NeuN labelling is superimposed on a dark-field image (grey). The superior cerebellar peduncle (SCP) is outlined in each image, and the approximate locations of the subnuclei have been plotted onto the middle image, based on the Franklin and Paxinos atlas^34^, and corresponding to a level of 1.54 mm behind the interaural plane. Within the lateral parabrachial (LPB) area, there is a very high density of labelling in the most dorsal aspect of the LPB, which largely corresponds to the internal lateral nucleus (il). Apart from this, there are only a few labelled fibres and boutons in other parts of the LPB. Note that this section is at a slightly more rostral level than the one shown in Fig 2, and the bundle of fibres travelling on the dorsolateral aspect of the brainstem to enter PBil is therefore not seen. Abbreviations for other subnuclei of the LPB: cl, centrolateral; dl, dorsolateral; el, external lateral; vl, ventral lateral. Scale bar = 200 μm.

Axonal labelling in the rostral brainstem and thalamus was best seen in mice that had survived 8 weeks after spinal injection of AAV.Cre^ON^.tdTom (Fig 4). In these animals, labelled axons were seen at a moderate density throughout the rostrocaudal extent of the PAG on both sides. Only a few sparsely distributed fibres were present in the superior colliculus (not shown). Within the thalamus, labelled axons were seen in the posterior triangular (PoT) nucleus and these were more numerous on the side contralateral to the injection site (Fig 4B,D). Apart from this, thalamic labelling was restricted to the medial part, in particular the mediodorsal (MD) nucleus (Fig 4A,C). There was essentially no labelling in two other thalamic nuclei that are known to receive input from the ALS: posterior (Po) and ventral posterolateral (VPL) nuclei. A few labelled axons were seen in the hypothalamus.

**Fig 4.**
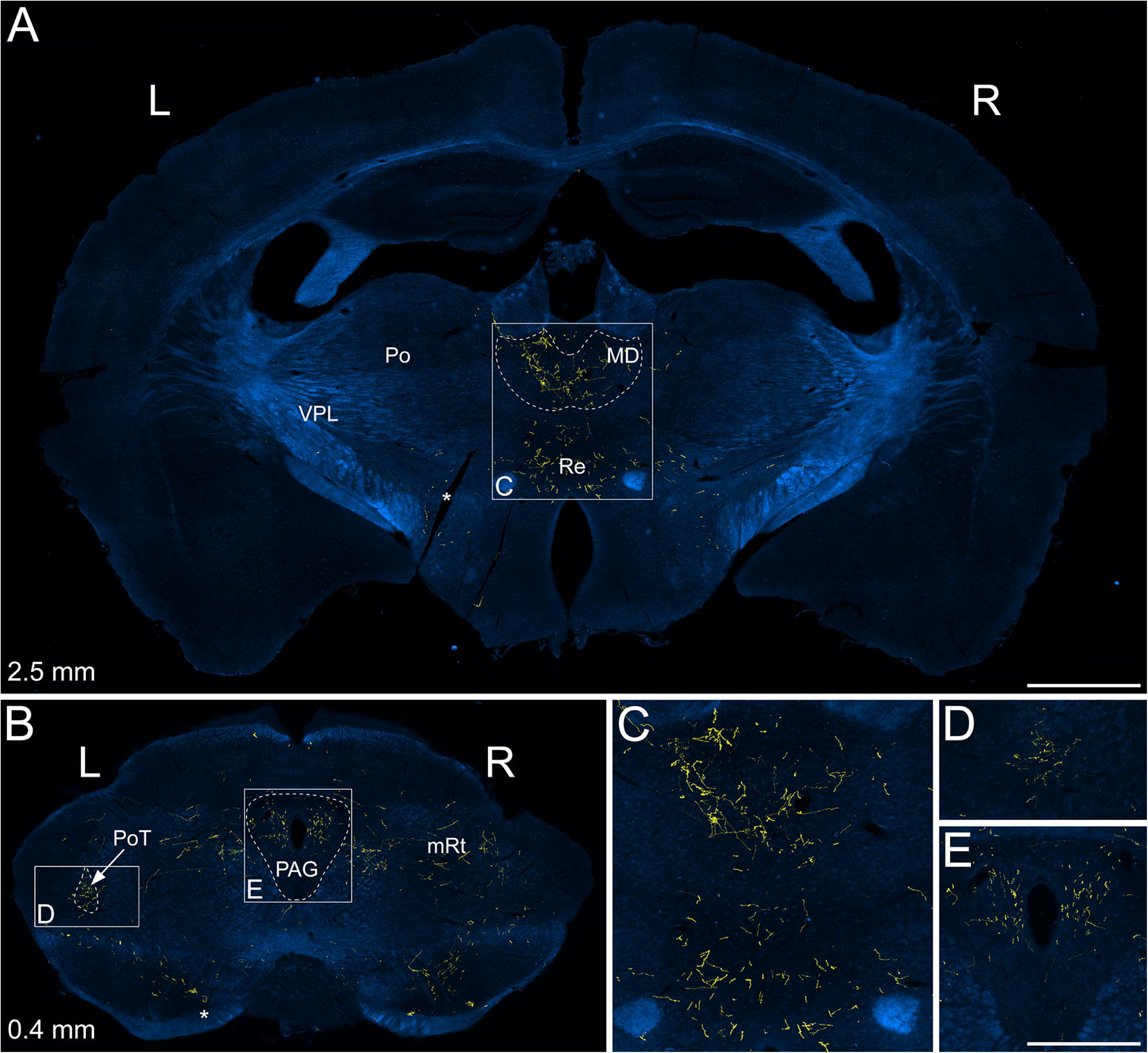
TdTomato labelling in the midbrain and forebrain of a Sst^Cre^ mouse that had received intraspinal injection of AAV.Cre^ON^.tdTomato into the L3, L4 and L5 segments on the right side ∼8 weeks previously (#6 in Table 1). TdTomato (yellow) is superimposed on dark-field images (blue) and approximate interaural levels are indicated^34^. **A**: tdTomato-positive axons are seen within the medial part of the thalamus at this level. These are particularly numerous in the mediodorsal nucleus (MD), with some labelling also in the nucleus reuniens (Re). No tdTomato axons are seen within either the ventral posterolateral (VPL) or posterior (Po) nucleus of thalamus. **B**: TdTomato-labelled axons are seen within the PAG on both sides and in the posterior triangular nucleus of the thalamus (PoT). In PoT, axons are more numerous on the left side (i.e., contralateral to the injection site). Other labelled axons are scattered through parts of the midbrain reticular formation (mRt). Asterisks indicate a cut made on the left side that was used to orientate the sections. **C**, **D** and **E** show regions indicated by the boxes in **A** and **B** at higher magnification. Scale bars = 1 mm (**A**,**B**) and 500 μm (**C**-**E**).

Only very occasional labelled axons were observed in the cerebellum, and none were present in the gracile nuclei, indicating minimal expression of *Sst* by spinocerebellar or postsynaptic dorsal column pathways. Since axonal labelling in the brain was restricted to known targets of the ALS (NTS, various reticular nuclei, LPB, PAG, thalamus), this demonstrates that at least the vast majority of *Sst*-expressing lumbar projection cells belong to this system.

### Expression of *Sst* in retrogradely labelled lamina V neurons

We previously showed that the ALS4 cluster (which includes Sst-expressing cells) was mainly located in the lateral reticular part of lamina V, and that most of the retrogradely labelled cells seen when AAV.Cre^ON^.tdTomato was injected into the LPB were present in this region^27^. We therefore determined the proportion of projection neurons in the lateral part of lamina V that expressed *Sst*. We injected cholera toxin B subunit (CTb) bilaterally into the caudal ventrolateral medulla (CVLM) of two Phox2a::Cre;Ai9 mice and combined immunohistochemical detection of CTb and tdTomato with fluorescent *in situ* hybridisation (FISH) for *Sst* mRNA on 16 μm thick transverse cryostat sections. The CVLM was selected for CTb injections because this region contains the main ascending axon bundle belonging to *Sst*-expressing ALS neurons (Fig 2).

Numerous CTb-labelled cells were seen in the L4 segments of these animals, and these were located in several regions of the spinal cord, including lamina I, the deep dorsal horn (laminae III-VI), the lateral white matter, the LSN and the region around the central canal. Some of these cells were also labelled with tdTomato, indicating that they were derived from the Phox2a lineage. The distribution of cells that were labelled with CTb and/or tdTomato in the lateral part of lamina V (including its lateral reticulated portion) is shown in Fig 5. Many of these cells contained *Sst* mRNA, and consistent with our previous findings^27^, these were concentrated in the reticulated part of lamina V, which can be readily identified when sections are viewed with dark-field illumination^27^. We therefore restricted our analysis to this region, and found that the great majority (71%, 78%, mean 75%) of retrogradely labelled cells were *Sst*-positive (Table 2). Most of the retrogradely labelled cells in this region (89%, 83%) lacked tdTomato, whereas all tdTomato-positive cells were labelled with CTb. Expression of *Sst* did not appear to differ between the cells with or without tdTomato (Table 2).

**Fig 5.**
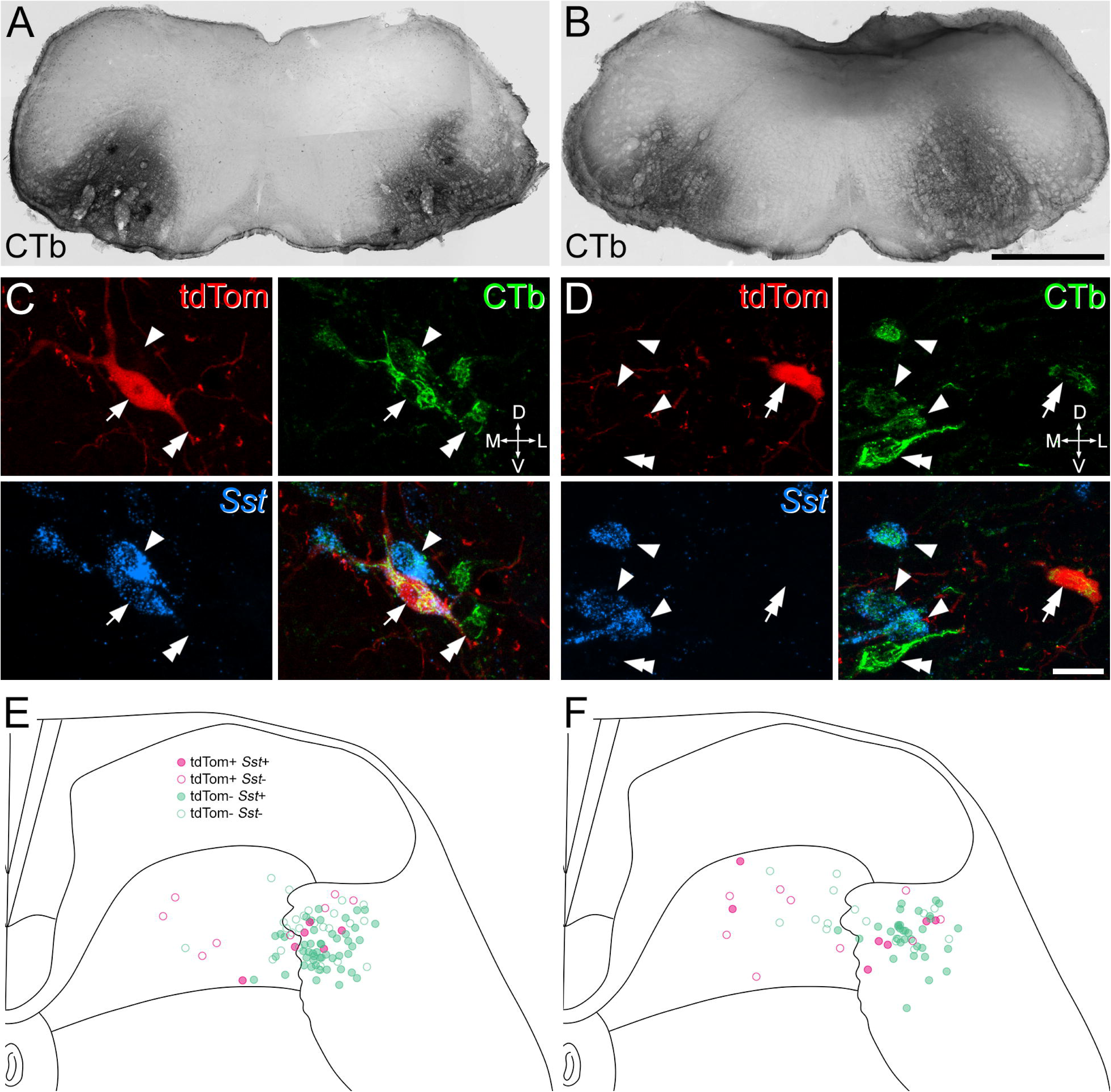
Expression of *Sst* mRNA in projection neurons in lamina V. **A**, **B**: CTb-labelling in the injection sites of the 2 Phox2a::Cre;Ai9 mice that were used for this analysis. Injections were targeted to the caudal ventrolateral medulla (CVLM) on both sides. **C**, **D**: examples of immunostaining for CTb (green) and tdTomato (red), and of fluorescent in situ hybridisation for *Sst* mRNA (blue) in each animal. In both cases, a region occupying the lateral reticulated part of lamina V is shown. The arrow in **C** shows a tdTomato+, CTb-labelled cell with *Sst* and the double arrow in **D** points to a tdTomato+, CTb-labelled cell that lacks *Sst*. Single and double arrowheads in both sets of images indicate tdTomato-negative, CTb-labelled cells that are positive (single arrowhead) or negative (double arrowhead) for *Sst*. **E**, **F**: Plots of the locations of all retrogradely labelled lamina V cells in each animal. Pink and green symbols represent tdTomato+ and tdTomato-negative cells, respectively. Filled symbols show cells with *Sst* mRNA and open symbols show cells that lacked *Sst* mRNA. Confocal scans in **C** and **D** are maximum intensity projections of 6 and 7 optical sections, respectively, at 1 μm z-separation. Orientation markers in **C**, **D**: D, dorsal; V, ventral; L, lateral; M, medial. Scale bars: **A**, **B** = 1 mm; **C**, **D** = 20 μm.

**Table 2.** The proportions of retrogradely labelled cells in the lateral part of lamina V with *Sst* mRNA. CTb, cholera toxin B subunit; tdTom, tdTomato

| Animal | All CTb+ | CTb+/tdTom- | CTb+/tdTom+ |
| --- | --- | --- | --- |
| 1 | 46/65<br>(70.7%) | 42/58<br>(72.4%) | 4/7<br>(57.1%) |
| 2 | 36/46<br>(78.3%) | 31/38<br>(81.6%) | 5/8<br>(62.5%) |

To provide an estimate of the total number of projection neurons in the lateral part of lamina V in the L4 segment, we re-analysed the scans using a variant of the disector method^35^, by including only those cells for which the bottom surface of the nucleus (stained with DAPI) was contained within the 16 μm cryostat section. We found 42 projection neurons in 20 sides (2.1 cells per section per side) in one animal and 22 from 14 sides (1.57 cells per section per side) in the other. This gives a mean value of 1.8 cells per 16 μm section on one side, and since we have previously reported that the L4 segment is 1.45 mm long^36^, we estimate that the total number of projection neurons in the lateral white matter (excluding the LSN) is ∼165 on each side, with around 75% of these (corresponding to ∼124 cells) expressing *Sst* and presumably belonging to the ALS4 population.

### Relation to other molecularly defined populations among the ALS

Our findings that *Sst*-expressing ALS cells are mainly located in the lateral reticulated part of lamina V^27^, and that their main target within the parabrachial area is the internal lateral (PBil) nucleus, are consistent with the findings of previous anterograde tracing studies in the rat, which showed that PBil is innervated by spinal projection neurons located in the deep grey matter, including cells in the lateral part of lamina V, but not by neurons in superficial laminae^6,7^. However, Choi et al^10^ reported that PBil also receives input from two ALS populations, defined by expression of Tacr1 and Gpr83. Although they had focused on ALS cells in the SDH that expressed these genes, the innervation of PBil suggests that some of the cells were located in deeper regions. In addition, we had found significant expression of *Tacr1* and some expression of *Gpr83* in cells belonging to ALS4 in our snRNAseq analysis^27^ (Fig S3A). We therefore looked for evidence of expression of Tacr1 and Gpr83 in lateral lamina V cells by injecting CTb into the left CVLM of two Tacr1^CreERT2^;Ai9 and two Gpr83^CreERT2^;Ai9 mice^10^. In these mice, cells that express Tacr1 or Gpr83 at the time of administration of tamoxifen are labelled with tdTomato. Since most axons belonging to *Sst*-expressing ALS neurons ascend on the ipsilateral side of the medulla, we restricted our analysis to lateral lamina V cells that were ipsilateral to the CTb injection, and quantified tdTomato labelling in cells that contained CTb. To provide further evidence for Tacr1 expression in the Tacr1^CreERT2^;Ai9 mice, we also immunostained these sections for the neurokinin 1 receptor (NK1r), the product of the Tacr1 gene. Examples of immunostaining, together with plots of the cells and injection sites, are illustrated in Figs 6 and S3B and quantitative results from the Tacr1^CreERT2^;Ai9 experiments are shown in Table 3. In the 2 Tacr1^CreERT2^;Ai9 mice, we found that 71% and 39% of CTb-labelled cells in this region contained tdTomato, while 61% and 60% were NK1r-immunoreactive. As expected, there was considerable overlap between tdTomato and NK1r expression, with 80% of tdTomato-positive cells staining for NK1r, and 71% of NK1r-immunoreactive cells expressing tdTomato. In tissue from the 2 Gpr83^CreERT2^;Ai9 we found that 20% and 59% of CTb-labelled cells in the lateral part of lamina V on the ipsilateral side were tdTomato-positive. This variability probably reflected inter-animal differences in the level of expression of tdTomato in these crosses. However, overall our results show that some projection neurons in the lateral reticulated part of lamina V are captured in each case, and therefore presumably express Tacr1 or Gpr83.

**Fig 6.**
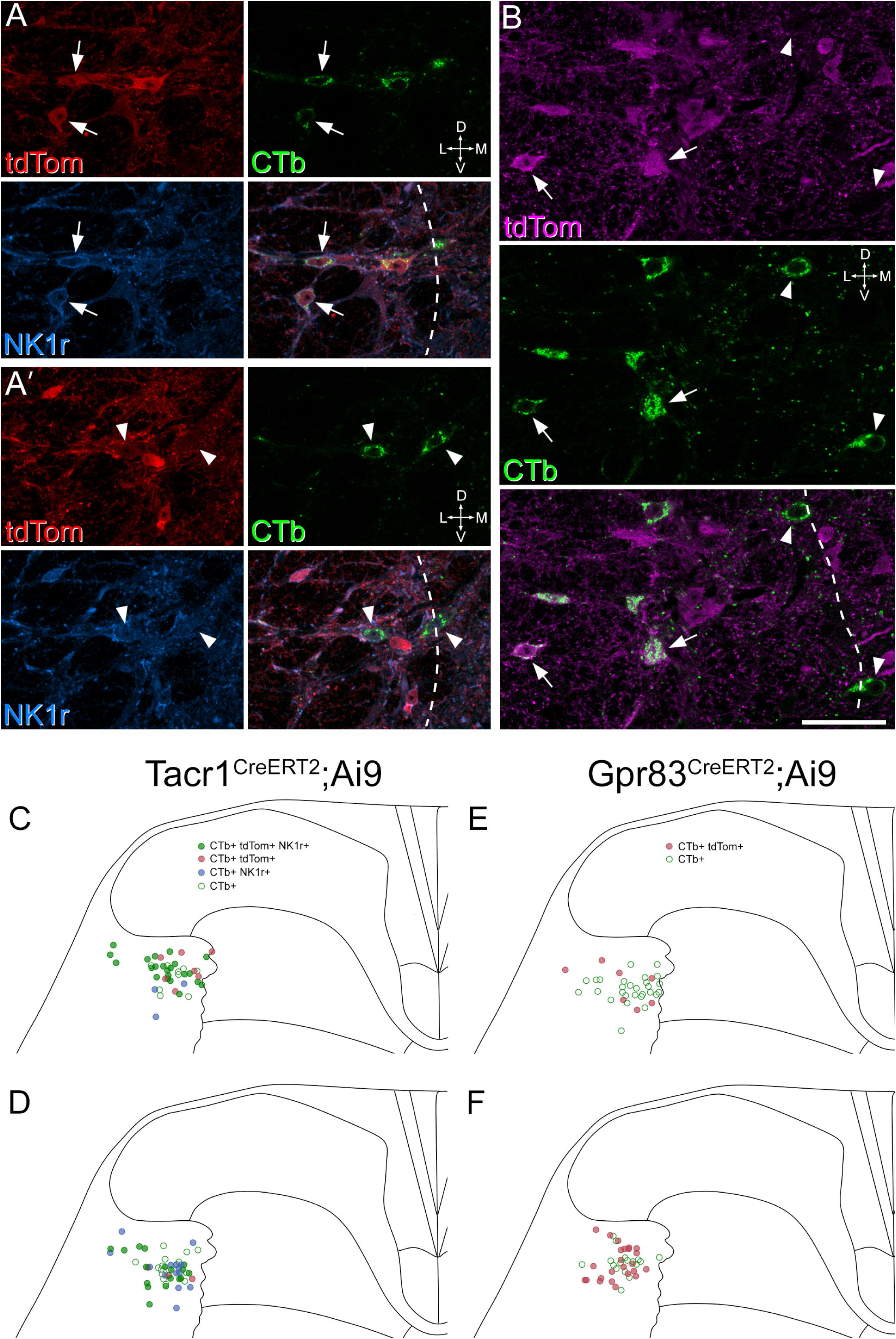
Expression of tdTomato and neurokinin 1 receptor (NK1r) in retrogradely labelled neurons in Tacr1^CreERT2^;Ai9 and Gpr83^CreERT2^;Ai9 mice. **A** and **A’** show two different planes through a transverse section from a Tacr1^CreERT2^;Ai9 mouse immunoreacted for tdTomato (tdTom, red), CTb (green) and NK1r (blue). Two CTb-labelled cells that contain tdTomato and express the NK1r are shown with arrows in **A**, and two CTb-labelled cells that lack tdTomato and are not NK1r-immunoreactive are shown with arrowheads in **A’**. **B** shows part of a section from a Gpr83^CreERT2^;Ai9 mouse immunostained to reveal tdTomato (tdTom, magenta) and CTb (green). Several CTb-labelled cells are visible in this field. Two of those that contain tdTomato are indicated with arrows, and two that lack tdTomato are shown with arrowheads. In both **A** and **B**, the dashed lines indicate the lateral border of the grey matter. **C**-**F** show plots of the distribution of retrogradely labelled cells in the lateral reticulated part of lamina V in two Tacr1^CreERT2^;Ai9 mice (**C**, **D**) and two Gpr83^CreERT2^;Ai9 mice (**E**, **F**). In **C** and **D**, green circles show retrogradely labelled cells with tdTomato and NK1r, brown circles cells with tdTomato that were NK1r-negative, blue circles those without tdTomato that were NK1r-immunoreactive and empty green circles those that lacked tdTomato and NK1r-immunoreactivity. In **E** and **F**, brown circles represent retrogradely labelled cells with tdTomato, and empty green circles indicate cells that lack tdTomato. Orientation markers in **A**, **B**: D, dorsal; V, ventral; L, lateral; M, medial. Scale bar for **A** and **B** = 50 μm.

**Table 3.** Expression of tdTomato and neurokinin 1 receptor (NK1r) in neurons retrogradely labelled with cholera toxin B subunit (CTb) in Tacr1^CreERT2^;Ai9 mice. The 2^nd^-5^th^ columns show the proportion (and percentage) of retrogradely labelled cells with each type of immunoreactivity. The last two columns show the proportion of tdTomato-positive cells that were NK1r-immunoreactive (or vice versa) among CTb-labelled cells in lateral lamina V.

| Animal | CTb only | CTb+/NK1r+/tdTom- | CTb+/NK1r-/tdTom+ | CTb+/NK1r+/tdTom+ | % tdTom with NK1r | %NK1r with tdTom |
| --- | --- | --- | --- | --- | --- | --- |
| 1 | 9<br>(22%) | 3<br>(7%) | 7<br>(17%) | 22<br>(54%) | 76% | 88% |
| 2 | 18<br>(35%) | 14<br>(27%) | 3<br>(6%) | 17<br>(33%) | 85% | 55% |
| **Mean** | **28%** | **17%** | **11%** | **43%** | **80%** | **71%** |

Another population that had been identified among ALS cells is characterised by expression of Tac1^37^, the gene that encodes preprotachykinin A, the precursor for substance P. Tac1 is widely expressed among Phox2a-derived projection neurons, including many of those in ALS4^27^ (Fig S4A). Consistent with this, Huang et al reported that Tac1 projection neurons were located in various locations, apparently including the lateral reticulated area of lamina V^37^. They identify the main parabrachial target as the superior lateral nucleus, but this has a similar location to PBil. To test this, we reviewed sections from a previous study in which we had anterogradely labelled Tac1-expressing projection neurons^21^, and confirmed that axons of these cells terminate bilaterally in PBil (Fig S4B). This difference likely results from variations in the nomenclature of LPB subnuclei.

Together, these findings support the suggestion that some cells in each of these molecular classes (defined by expression of *Tacr1*, *Gpr83* or *Tac1*) belong to the ALS4 population, and that these contribute to the innervation of PBil.

### Retrograde transport of AAVs to target *Sst*-expressing ALS neurons

We had previously shown that a unilateral injection of AAV coding for Cre-dependent tdTomato into the LPB in Sst^Cre^ mice resulted in retrograde labelling of cells in lateral lamina V on both sides. This symmetrical arrangement presumably occurs because many of the ascending axons of these cells, having passed through the ipsilateral LPB, cross the midline in the SMV to reach the contralateral LPB (Fig 2). We predicted that if this was the case, injection of retrogradely transported virus into the CVLM on one side should result in mainly ipsilateral labelling of ALS neurons. To test this, we injected AAV2r.Cre^ON^.GFP into the left CVLM of 2 Sst^Cre^ mice. The choice of this serotype was based on a recent report by Browne et al that AAV2r selectively targets ALS neurons in lateral lamina V^38^. As expected, we found that retrogradely labelled neurons were largely restricted to the lateral part of lamina V, and these were far more numerous on the side ipsilateral to the CVLM injection (Figs 7 and S5, Table 4). Across the two mice, the mean number of cells in lateral lamina V on the ipsilateral side was 1.9 per 60 μm section (corresponding to 46 cells in the L4 segment^36^), with 0.36 cells per section on the contralateral side. Cells in the ipsilateral lamina V therefore outnumbered those in the contralateral lamina V by a ratio of 5.5:1.

**Fig 7.**
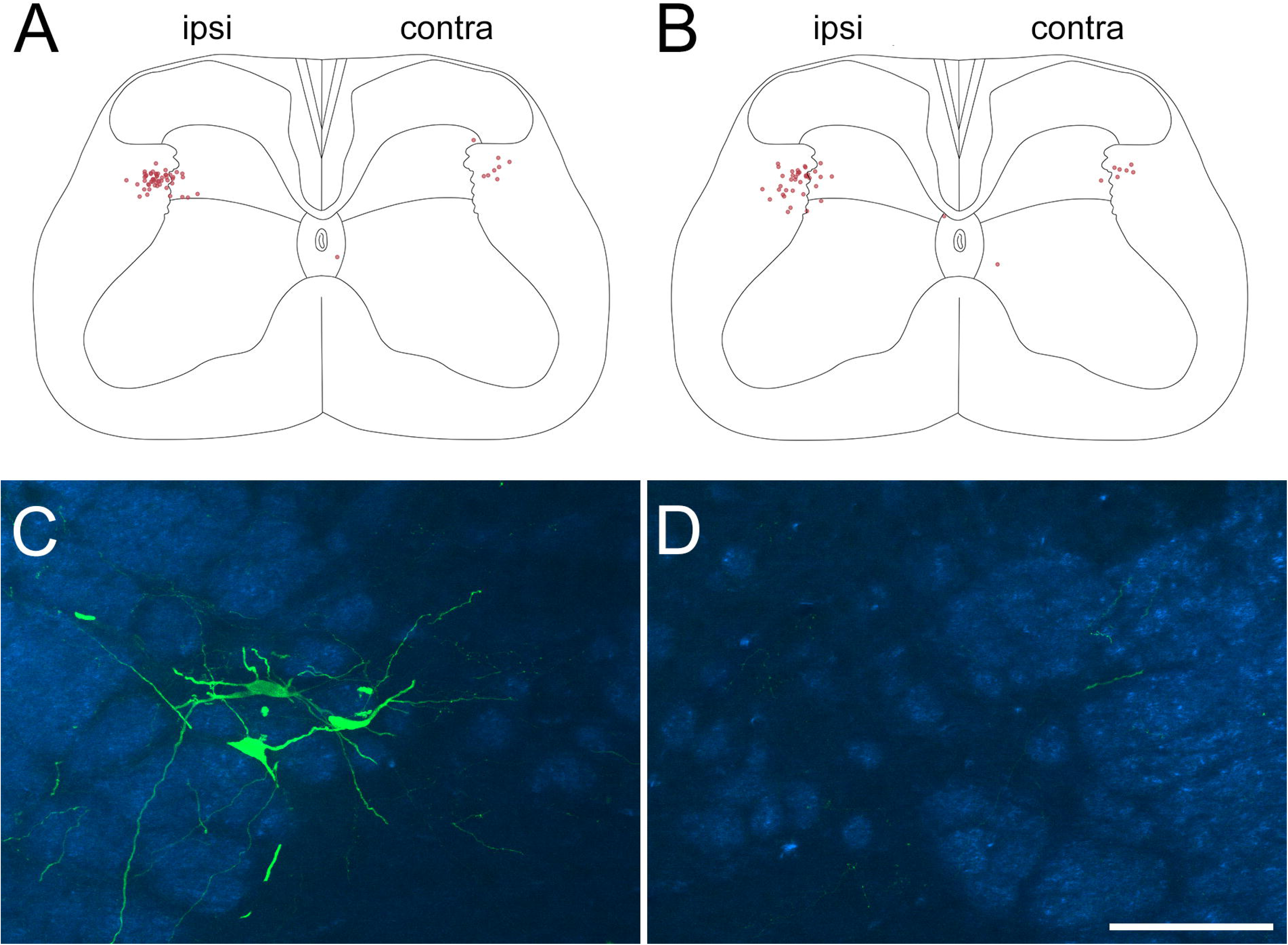
Retrograde labelling of ALS neurons following injection of AAV2r.Cre^ON^.GFP into the left CVLM of two Sst^Cre^ mice. **A**, **B** show plots of all of the retrogradely labelled cells identified in the L4 segment in each case. All but three of the labelled cells are located in the lateral part of lamina V, with the remaining three in or near lamina X, which surrounds the central canal. The retrogradely labelled cells are mainly present on the left side, ipsilateral (ipsi) to the CVLM injection, with many fewer cells on the contralateral (contra) side. **C**, **D** show the lateral part of lamina V on the ipsilateral (**C**) and contralateral (**D**) sides of one of the sections from the animal illustrated in **B**. GFP labelling is shown in green, superimposed on a dark-field image (blue). Note the presence of retrogradely labelled cells on the ipsilateral, but not the contralateral, side in this section. Scale bar = 100 μm. Injection sites for these two mice are shown in Fig S4.

**Table 4.** Retrograde labelling of lateral lamina V cells following unilateral injection of AAV2r.Cre^ON^.GFP into the CVLM of Sst^Cre^ mice.

| Animal | Sections analysed | Cells in ipsi lamina V | Cells in contra lamina V | Cells in ipsi lamina V per section | Cells in contra lamina V per section | Ratio of ipsi lamina V : contra lamina V |
| --- | --- | --- | --- | --- | --- | --- |
| 1 | 27 | 54 | 8 | 2 | 0.3 | 6.75 |
| 2 | 21 | 38 | 7 | 1.81 | 0.33 | 5.43 |
| **Mean** |  |  |  | **1.9** | **0.31** | **6.09** |

Finally, to provide further information on spinal cord collaterals of the *Sst*-expressing ALS cells, we targeted these neurons by injecting AAV.Flp into the CVLM on one side, and AAV.Cre^ON^/Flp^ON^.GFP^39^ into the lumbar spinal cord on the same side, in two Sst^Cre^ mice. Cell body labelling with GFP was seen in the injected segments, and this was largely restricted to neurons in the lateral part of lamina V (Fig 8A). In addition, a few labelled axons were seen in lamina I close to the site of the spinal injection (Fig. 8A inset). In segments rostral to the spinal injection sites, a bundle of ascending axons was present in the ipsilateral lateral funiculus, and collateral branches were seen entering the lateral part of lamina V and extending towards the central canal (Fig 8B). In one of the mice, which survived for 29 days after the operation, these collaterals were seen as far rostrally as the C7 segment, while in the other (which survived for 17 days) they could be identified as far as the T11 segment. This confirms that the collateral branches seen in these locations following intraspinal injection of AAV.Cre^ON^.tdTomato in the Sst^Cre^ mice (Figs 1, S1) originated from ascending axons of the *Sst*-expressing ALS cells.

**Fig 8.**
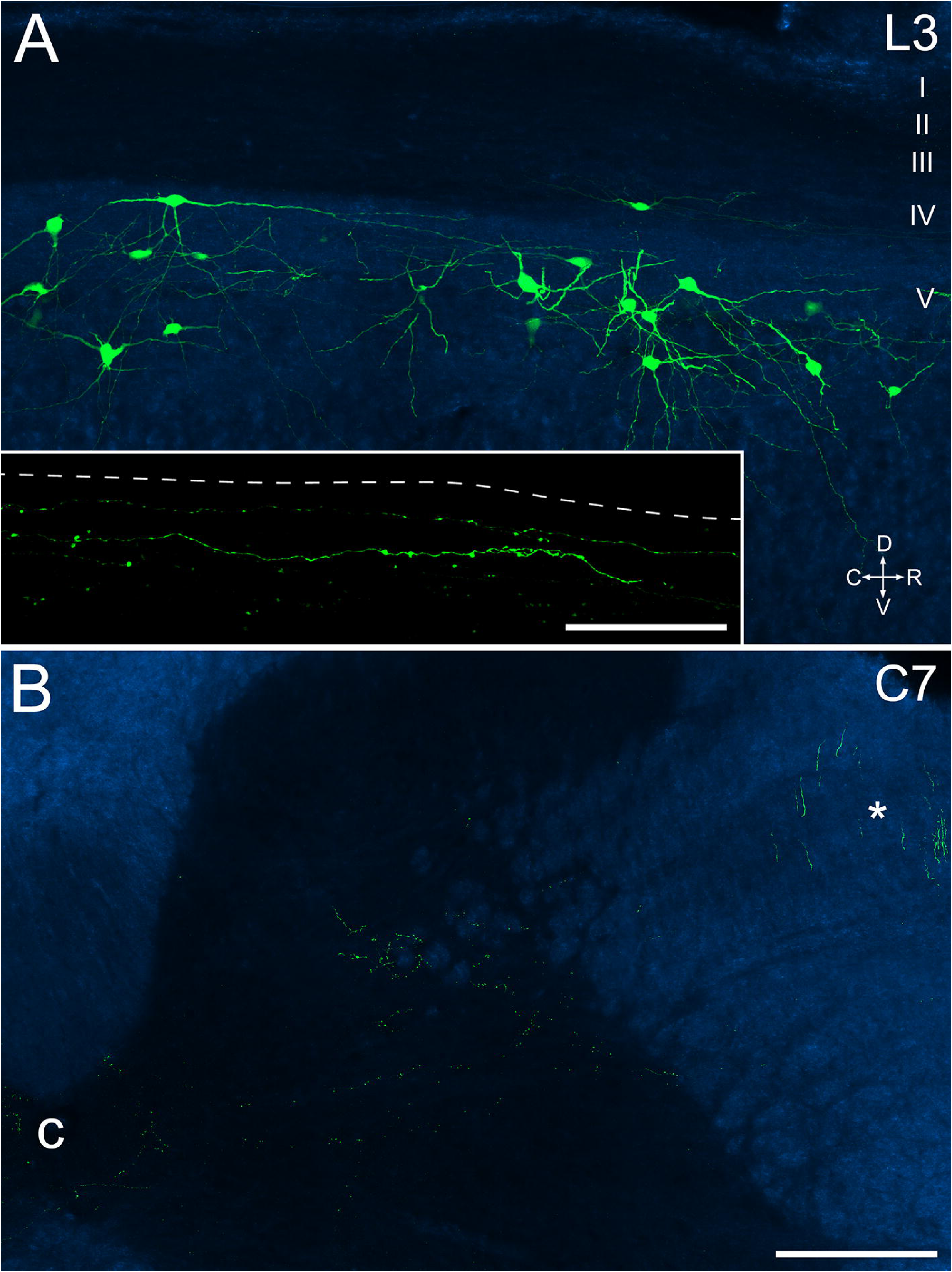
Retrograde labelling of Sst-expressing ALS neurons following injection of AAV.Flp into the CVLM on one side and injection of AAV.Cre^ON^/Flp^ON^.GFP into the lumbar enlargement on the same side in a Sst^Cre^ mouse. The injections were performed 29 days before perfusion fixation. **A**: A sagittal section through the L3 segment shows several cells labelled with GFP (green) superimposed on a dark-field image (blue). Approximate laminar levels are shown on the right. Orientation marker: D, dorsal; V, ventral; C, caudal; R, rostral. Inset shows part of the superficial dorsal horn from the same section scanned at a higher magnification to show GFP-labelled axons. The dashed line shows the position of the grey/white matter border. **B**: A transverse section from the C7 spinal segment of the same animal. Again, GFP labelling (green) is superimposed on a dark-field image (blue). This image shows the presence of GFP-labelled axons ascending in the lateral funiculus (*). GFP-labelled boutons belonging to collateral axons can be seen in the grey matter, and these are present in the lateral part of lamina V and in the area around the central canal. Collateral axons in the superficial dorsal horn are not seen at this segmental level. c, central canal. Scale bar for **A** and **B** = 200 μm; Scale bar for inset in **A** = 50 μm.

## DISCUSSION

The main findings of this study are that somatostatin-expressing neurons account for ∼75% of ALS cells in lateral lamina V, and project mainly through the ipsilateral lateral funiculus to the PBil on both sides, while giving off collateral branches at multiple spinal levels, as well as to several brainstem areas. Thalamic projections from somatostatin-expressing ALS cells are present in PoT, but are otherwise restricted to the medial nuclei, especially MD.

### Somatatostatin as a marker for ALS4 neurons

We previously reported that *Sst* was among the top differentially expressed genes within the ALS4 cluster^27^. *Sst* was not detected in all ALS4 neurons, and it is occasionally expressed in cells belonging to other clusters. Nonetheless, our finding that Cre-expressing neurons in Sst^Cre^ mice retrogradely labelled from the LPB^27^ or CVLM (present study) are largely restricted to the lateral part of lamina V suggest that the Sst^Cre^ mouse provides a reliable way of targeting ALS4 cells and that these have a very distinctive distribution within the spinal cord.

Browne et al^38^ have recently demonstrated that viruses of the AAV2retro serotype injected into LPB selectively label ALS neurons in lateral lamina V, suggesting that they show a tropism for ALS4. However, the density of cells that they captured was relatively low: 0.5 cells per 50 μm, which is equivalent to 15 cells per side in the L4 segment. We confirmed that this serotype is able to retrogradely label ALS4 cells in lateral lamina V, by injecting AAV2r.Cre^ON^.GFP into Sst^Cre^ mice. However, we achieved ∼3-fold higher number of retrogradely labelled neurons, possibly because of more efficient labelling from the CVLM. By comparing results from these injections with the total number of *Sst*-containing projection neurons in lateral lamina V, we estimate that ∼37% of these cells were GFP-labelled, and this proportion would be slightly higher if any cells project exclusively via the contralateral CVLM. Although Browne et al describe the lateral lamina V cells that they captured as being a small population^38^, we estimate that there are ∼120 *Sst*-positive ALS cells in lateral lamina V in L4. We previously reported that there are around 240 lamina I ALS neurons in this segment^40^, and lamina I contains at least 3 different transcriptomic classes of projection neuron^27^. It is therefore likely that the ALS4 population is of a comparable size to (or even larger than) other ALS populations.

Although our transcriptomic classification was based on those ALS cells that are derived from the Phox2a lineage^27^, we found that only around 15% of the *Sst*+ ALS cells in two Phox2a::Cre;Ai9 mice were labelled with tdTomato. While it is possible that *Sst* is expressed in ALS neurons that are not derived from this lineage, a more likely explanation is that lack of tdTomato in many of these cells reflects incomplete capture of Phox2a-derived cells in this BAC transgenic line. Consistent with this, Roome et al^9^ reported that only a minority of lamina V cells with *Phox2a* mRNA were tdTomato-positive in Phox2a::Cre;Ai14 mice at E16.5, and we have found considerable inter-animal variation in the extent to which deep dorsal horn ALS neurons were labelled in Phox2a::Cre;Ai9 mice^41^.

Other molecular markers used to define ALS populations in recent studies include the genes coding for Tacr1 and Gpr83^10^, and for Tac1^37^. However, it appears that all three of these are expressed by at least some of the ALS cells in the lateral reticulated part of lamina V, as well as by ALS cells in other locations, suggesting that they are not restricted to single populations. In contrast, the highly specific laminar location of the *Sst*-expressing cells supports the relatively selective association of *Sst* with a single population of ALS cells.

### Response properties of ALS4 neurons

Numerous studies of the receptive field properties of ALS projection neurons located in lateral lamina V have been carried out in a variety of species^12,42^. These cells are generally described as having wide dynamic range (WDR) receptive fields, responding weakly to innocuous cutaneous stimuli and strongly to noxious mechanical or thermal stimulation of the skin^43–45^, with some cells also being activated by visceral stimulation^46^, or application of pruritogens to the skin^44^. In addition, Menetrey et al^47^ reported that projection neurons in lateral lamina V upregulated the activity-dependent marker Fos following subcutaneous injection of complete Freund’s adjuvant or periarticular injection of urate crystals, suggesting that these cells have a role in inflammatory pain states.

The ALS4 cells are likely to receive only a sparse direct synaptic input from nociceptors, as these terminate mainly in the SDH, and their nociceptive input is likely to be mainly transmitted indirectly through excitatory interneurons^41^. One potential population of excitatory interneurons that could provide this input consists of GRPR-expressing cells in laminae I-II. These largely correspond to vertical cells, and have axons that arborise in lateral lamina V^48^. We have shown that chemogenetic activation of the GRPR cells evokes behaviours that appear to reflect both pain and itch^48^, and results in Fos expression in many cells in lateral lamina V, including projection neurons^49^. The GRPR cells may therefore provide both nociceptive and pruritoceptive input to ALS4 cells.

### The course of ALS4 axons within the spinal cord

One highly unusual feature of the projection from the somatostatin-expressing ALS cells was that the main bundle of ascending axons was located on the ipsilateral side in the spinal cord. We initially observed this when examining anterograde transport of AAV.Cre^ON^.tdTomato injected into the lumbar spinal cord of Sst^Cre^ mice, and subsequently confirmed it by showing that most retrogradely labelled ALS cells were located ipsilateral to an injection of AAV2r.Cre^ON^.GFP into the CVLM in these mice. In contrast, retrograde labelling of these cells with AAV.Cre^ON^.tdTomato injected into the LPB of Sst^Cre^ mice resulted in similar numbers on each side^27^, and Browne et al reported a symmetrical arrangement of cells in the lateral reticulated part of lamina V when AAV2r.GFP was injected into the LPB^38^. This suggests that having ascended on the ipsilateral side and entered the LPB on that side, axons of many of these cells cross the midline in the SMV, to enter the contralateral LPB.

This pattern is very different from that of most ALS neurons, which have axons that cross the midline close to the level of the cell body^11^ and ascend in the contralateral white matter^3,11,50^. Despite this difference, the course of the ascending fibres seems to mirror that seen by Chen et al^11^, who performed anterograde viral tracing from lumbar spinal cord (as shown in their Fig 2E,F). The functional significance of a mainly ipsilateral projection is not known, but if the same arrangement is present in the human spinal cord this would have implications for the interpretation of pain relief following anterolateral cordotomy. This procedure is performed on the side contralateral to the main site of pain, typically in patients with terminal malignancies, and results in effective pain relief on the side contralateral to the cordotomy, at least in the short term^51^. If ALS cells in lateral lamina V in humans also project through the ipsilateral side of the cord, then those located on the “painful” side would be spared by this procedure, suggesting that on their own, they are not essential for pain perception. However, in many cases the initial pain relief following cordotomy diminishes over time^51^, and an intact pathway from these ipsilateral ALS neurons may contribute to this phenomenon. In addition, some cordotomy patients report “mirror pain”, in which pain is felt on the same location as the original pain, but on the opposite side of the body^51^, and ipsilaterally projecting ALS cells may also be involved in this^11^.

Another unexpected feature was the presence of collateral axons that were apparently given off at all spinal levels rostral to the cell body. There has been considerable recent interest in collateral axons arising from spinal projection neurons, but this has mainly focused on those originating from cells in lamina I, and those given off close to the cell body^52,53^. Browne et al^38^ had combined optogenetic activation of ALS cells in lateral lamina V with detection of phosphorylated extracellular signal-regulated kinases to identify areas that received input from collateral axons, and found that these included the superficial and deep laminae of the dorsal horn, and the area around the central canal. Taken together with our anatomical findings, this suggests that there are two distinct patterns: collaterals that enter the SDH are given off within a few segments of the cell body, whereas those that innervate the deep dorsal horn and lamina X arise at multiple segmental levels. Browne et al proposed that the major targets of the local collaterals were inhibitory interneurons, and suggested that this could serve as a mechanism for suppressing activity in other cells^38^. Interestingly, these 3 target regions (SDH, lamina V and lamina X) all contain cell bodies of other ALS cells, and the collaterals may therefore influence other components of the ALS. However, it is not yet known whether the collaterals exert and inhibitory or facilitatory effect at each of these locations.

### Brain projections of ALS4 cells

Within the brainstem, the projection pattern of *Sst*-expressing ALS cells was generally similar to that described for other components of the ALS, with axons targeting the NTS, various reticular nuclei, the LPB and PAG. Consistent with previous studies that have traced projections from lateral lamina V^6,7^, we found that within the LPB complex, these cells projected mainly to PBil, and this highly restricted distribution supports the view that the Sst-expressing ALS cells represent a distinct functional population. Since projections traced in Tacr1^CreERT2^, Gpr83^CreERT2^ and Tac1^Cre^ mice also include this region, and ALS cells expressing these genes are present in the lateral part of lamina V, it is likely that these cells at least partly account for the PBil projections seen in these genotypes^10,37^.

Thalamic projections were restricted to the PoT and medial thalamic nuclei (in particular MD and nucleus reuniens). Gauriau and Bernard found that following injections of anterograde tracer into the lateral part of lamina V in rat cervical cord the main thalamic target was the centrolateral (CL) nucleus, which surrounds MD^5^. This difference may reflect a species or segmental difference, but more likely results from differences in approach, as theirs depended on spatial rather molecular specificity. Neurons in PBil project to the paracentral (PC) nucleus, and both PC and MD thalamus in turn project to the prefrontal cortex^54,55^, while the PoT thalamic nucleus targets the insular and second somatosensory cortices^56^. Taken together with the lack of input to VPL (which projects to primary somatosensory cortex), our findings suggest that activity in the ALS4 population will influence prefrontal, insular and S2 regions. It is therefore likely that these cells contribute to the motivation-affective, rather than sensory-discriminative, dimensions of pain.

## METHODS

### Animals

All experiments were approved by the Ethical Review Process Applications Panel of the University of Glasgow, and were performed in accordance with the European Community directive 86/609/EC and the UK Animals (Scientific Procedures) Act 1986. The study was carried out in compliance with the ARRIVE guidelines.

We used five genetically modified mouse lines. Many experiments involved a line in which Cre recombinase is knocked into the somatostatin locus^57^ (Sst-IRES-Cre, also known as Sst^Cre^; Jax strain 013044). In addition, we used Phox2a::Cre^9^ mice as well as Tacr1^CreERT2^ and Gpr83^CreERT2^ mice^10^, in each case crossed with the Ai9 reporter line, in which Cre-mediated excision of a STOP cassette drives expression of tdTomato. The Tacr1^CreERT2^;Ai9 and Gpr83^CreERT2^;Ai9 mice received single intraperitoneal injections of tamoxifen (1.5 mg for Tacr1^CreERT2^;Ai9 mice, 2.5 mg for Gpr83^CreERT2^;Ai9 mice^10^) at between 21 and 27 days of age.

Mice of both sexes, aged 5 to 12 weeks and weighing between 16 and 31 g at the start of surgical procedures, were used in this study.

### Surgery

All surgical procedures involving intraspinal and/or intracranial injections were performed while the mice were anaesthetised with isoflurane (∼1.5%), and all of these animals received perioperative analgesia (buprenorphine 0.1 mg/kg and carprofen 10 mg/kg, s.c.). Perfusion fixation was performed while mice were deeply anaesthetised with pentobarbitone (20 mg i.p.). Details of viral vectors and injection volumes are provided in Table S1.

Intraspinal injection of viral vectors coding for Cre-dependent constructs was performed as described previously^49,58^ on 8 Sst^Cre^ mice (Table 1). In 4 cases, the injections were made in the L3 segment, while in 4 cases injections were made into each of the L3, L4 and L5 segments. These mice underwent perfusion fixation after a survival period of 4-8 weeks. Two Sst^Cre^ mice (one of each sex) received injections of AAV2r.Cre^ON^.GFP targeted on the CVLM on the left side, and were fixed by perfusion 15 days later. Two Sst^Cre^ mice (one of each sex) were injected with AAV.Flp targeted to the left CVLM and AAV.Cre^ON^/Flp^ON^.GFP into the L3 and L5 spinal segments on the same side. They were perfused with fixative 17 or 29 days later. Two male Phox2a::Cre;Ai9 mice were injected with 300 nl of 1% CTb into the CVLM on both sides, and were perfused with fixative 2 days later. Two male Tacr1^CreERT2^;Ai9 and two female Gpr83^CreERT2^;Ai9 mice received injections of 300 nl of 1% CTb into the left CVLM and were perfused 2 or 3 days later. All injections into the brain were made using stereotaxic co-ordinates, as described previously^59^.

### Immunohistochemistry and image analysis

Multiple-labelling immunofluorescence reactions were performed as described previously^48,49^ on 60 μm thick transverse or parasagittal sections of spinal cord, or on 50 μm thick coronal sections of brain, which had been cut with a vibrating blade microtome (Leica VT1200 or VT1000) or a cryostat (Leica CM1950). Reactions were performed on free-floating sections. The sources and concentrations of antibodies used are listed in Table S2. Sections were incubated for 1-3 days at 4°C in primary antibodies diluted in phosphate buffered saline (PBS) that contained 0.3M NaCl, 0.3% Triton X-100 and 5% normal donkey serum, and then for 2-18 hours in appropriate species-specific secondary antibodies that were raised in donkey and conjugated to Alexa 488 or Rhodamine Red (from Jackson Immunoresearch, West Grove, PA) or to Alexa 555-plus (Thermo Fisher Scientific). All secondary antibodies were used at 1:500 (in the same diluent), apart from those conjugated to Rhodamine Red, which were diluted to 1:100. Following the immunohistochemical reaction, sections were mounted in anti-fade medium and stored at −20°C. Sections were scanned with Zeiss 710 LSM confocal microscope (Argon multi-line, 405 nm diode, 561 nm solid state and 633 nm HeNe lasers), using 10×, 40× or 63× oil-immersion objectives (numerical apertures of 0.3, 1.3 and 1.4, respectively). In all cases, the aperture was set to 1 Airy unit or less.

For the analysis of data from the Tacr1^CreERT2^;Ai9 and Gpr83^CreERT2^;Ai9 mice, we examined 5 transverse sections from the L4 segment of each animal. Only cells in the lateral reticulated part of lamina V on the side ipsilateral to the brain injection were examined. All CTb-labelled cells that were visible in this region were identified, and the presence of tdTomato and (for Tacr1^CreERT2^;Ai9 mice) NK1r immunoreactivity was noted.

For experiments in which AAV2r.Cre^ON^.GFP was injected into the CVLM of Sst^Cre^ mice, we examined 21 or 27 transverse sections from the L4 segment. GFP-labelled neurons were counted if they were entirely contained within the section, or if part of the soma was present at the top surface, but were excluded if part of the soma was present at the bottom of the section (i.e. if part of the soma appeared on the last scan in the z-series).

### Fluorescent *in situ* hybridisation and immunohistochemistry

Combined fluorescence *in situ* hybridisation (FISH) and immunohistochemistry was performed on sections from the L4 segments of the two Phox2a::Cre;Ai9 mice that had received injection of CTb into the CVLM, using a method described previously^27^. The tissue was cryoprotected, frozen, embedded in OCT, and then cut into 16 μm thick transverse sections on a cryostat (Leica CM1950). In situ hybridisation was carried out using a RNAscope multiplex fluorescent kit v2 according to the manufacturer’s instructions with probe against *Sst*, which was revealed with TSA Vivid 650. Slides were incubated overnight in antibodies against CTb and mCherry, which were revealed with secondary antibodies conjugated to Alexa 488 and Rhodamine Red (respectively), and stained with DAPI to reveal cell nuclei.

Confocal scans were acquired with the 10× and 40× lens, and analysed with Neurolucida for Confocal (MBF Bioscience, Williston, VT, USA). Since the CVLM injections were bilateral, we analysed tissue from both right and left sides of the spinal cord, with 20 or 14 sides being analysed from each mouse. Initial scans taken with the 10× lens included CTb and tdTomato immuno-fluorescence, combined with light transmitted through a dark field condensor. These scans were used to reveal the locations of labelled cells in relation to the grey/white matter border and Rexed’s laminae. Subsequent scans obtained with the 40× lens revealed nuclei (DAPI), CTb, tdTomato and *Sst* mRNA. We identified all neurons in the lateral half of lamina V and the lateral white matter (excluding the LSN) that were labelled with CTb and/or tdTomato, and for which at least part of the nucleus was present in the section. We then determined whether these cells were positive for *Sst* mRNA (based on the presence of 4 or more transcripts) and plotted their locations onto an outline of the spinal cord. To estimate the true number of lateral white matter projection neurons per section, we re-analysed the scans using a variant of the disector method^35^ and identified all of the CTb and/or tdTomato-labelled cells in this region for which the bottom surface of the nucleus lay within the (16 μm thick) cryostat section. This was achieved by examining z-series scanned through the entire thickness of the section, identifying the DAPI-stained nucleus in each optical section, and excluding any cells in which the nucleus was still present in the last optical section in the series.

## ACKNOWLEDGEMENTS

We thank Iain Plenderleith and Robert Kerr for expert technical assistance, Dr Artur Kania for the Phox2a::Cre mice and Dr David Ginty for the Tacr1CreERT2 and Gpr83CreERT2 mice. Financial support from the Wellcome Trust (Grant number 219433/Z/19/Z) and the Medical Research Council (Grant number MR/S002987/1) is gratefully acknowledged.

## AUTHOR CONTRIBUTION

A.M.B and A.J.T. conceived the project and designed experiments; W.M., E.P., A.C.D., M.A.H., R.Q., M.Y. and M.G.-M. performed experiments; W.M. and M.A.H. analysed data; M.W. provided reagents; J.H. and A.J.T. obtained funding; W.M, M.G.-M., A.M.B. and A.J.T. wrote the paper; all of the authors provided feedback and contributed to the editing of the manuscript.

## DATA AVAILABILITY

The datasets generated and analysed during the current study are available from the corresponding author on reasonable request.

