## Supplemental information for "Anatomical characterisation of somatostatin-expressing neurons belonging to the anterolateral system"

**Supplementary Figures 1-5**

**Supplementary Tables 1-2**

**Figure S1**

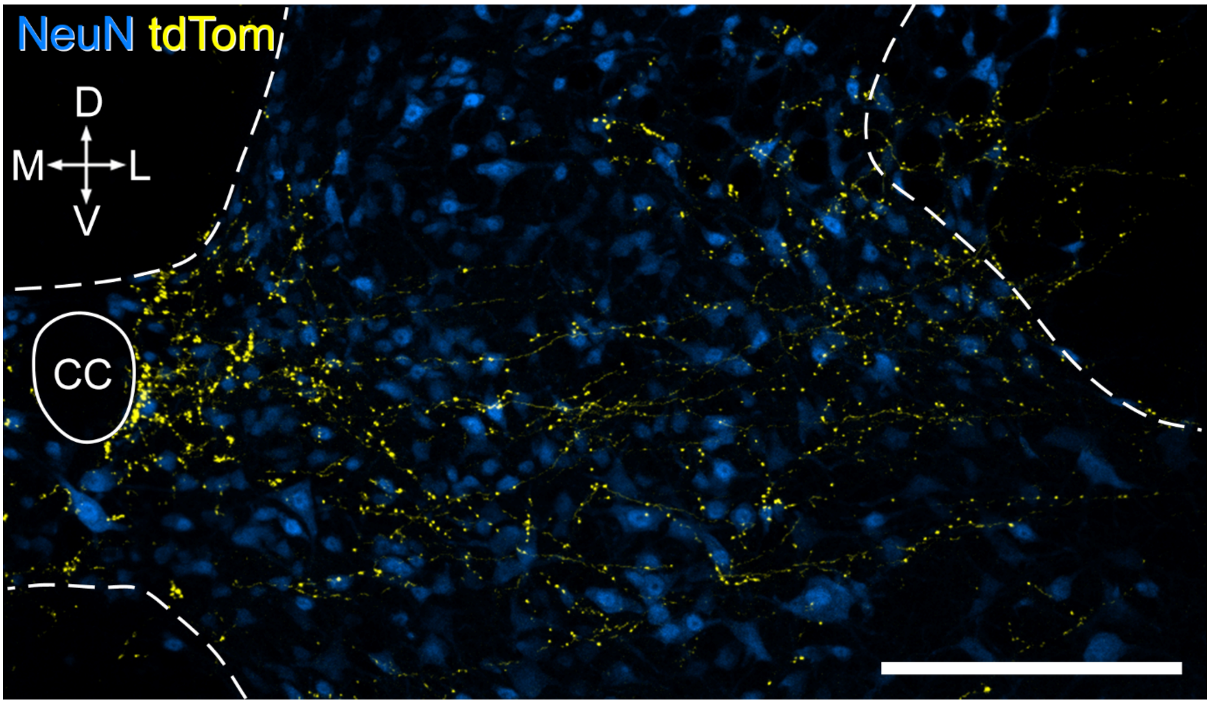


Axon collaterals in the cervical spinal cord. TdTomato labelling (yellow) on the ipsilateral side in the C7 spinal cord segment of a $Sst^{Cre}$ mouse that had received intraspinal injection of $AAV.Cre^{ON}$.tdTomato into the L3, L4 and L5 segments on the right side 5 weeks previously (#3 in Table 1). NeuN immunostaining is shown in blue. Note the presence of numerous tdTomato-labelled axons and boutons in the deep dorsal horn and the area around the central canal (cc). The dashed line represents the grey/white matter border. Orientation markers: D, dorsal; V, ventral; L, lateral; M, medial. Scale bar = 200 µm.

**Figure S2**

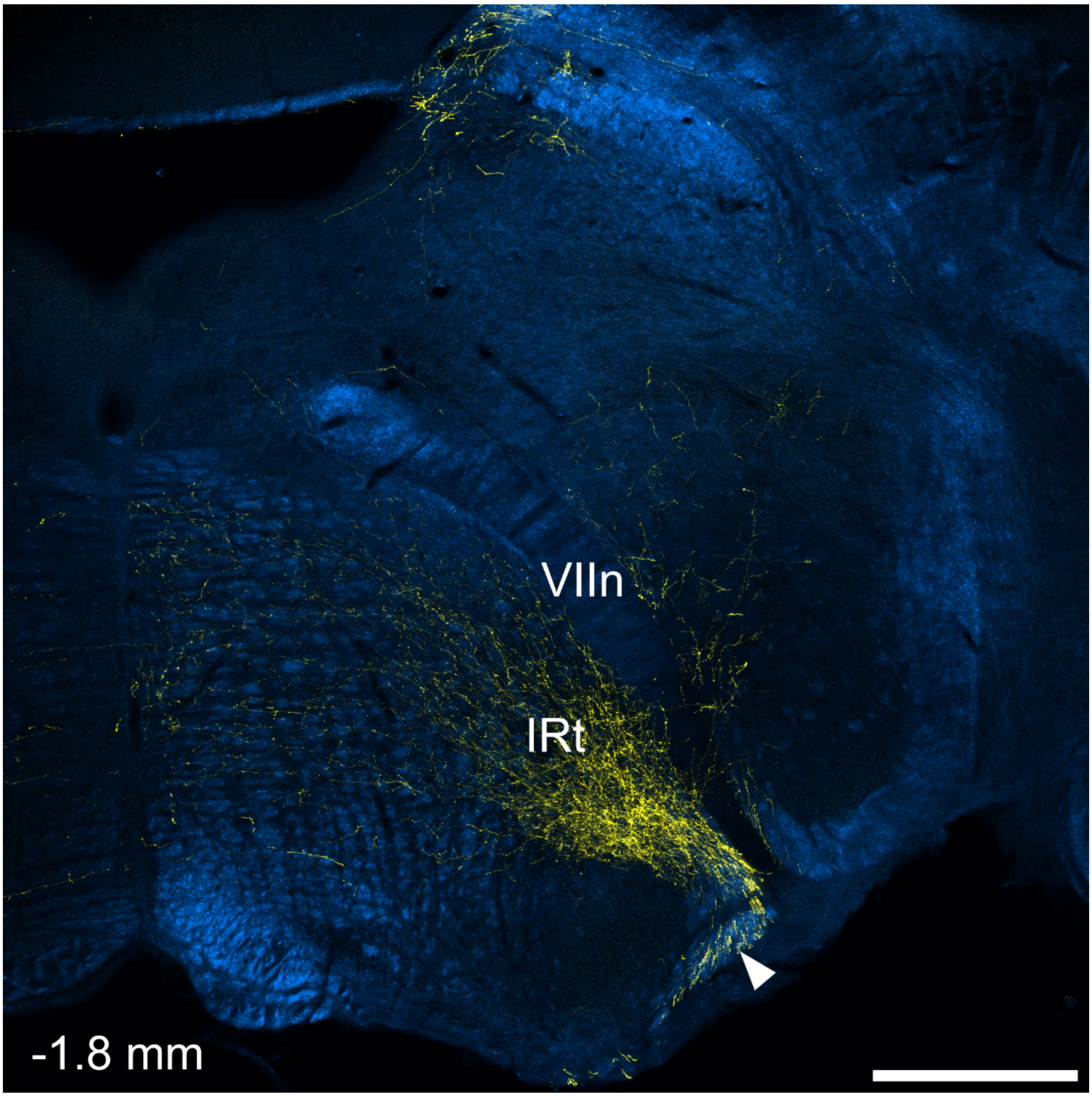


TdTomato labelling in a coronal section through the ipsilateral side of the brainstem of a $Sst^{Cre}$ mouse that had received an injection of AAV.$Cre^{ON}$.tdTomato into the L3 spinal segment on the right (#8 in Table 1). The section is located at approximately 1.8 mm caudal to the interaural line. The main ascending bundle of labelled fibres on the ventral surface of the brainstem is indicated with an arrowhead. This gives rise to a dense collection of axonal branches with boutons that lies on the ventromedial aspect of the facial nerve (VIIn) and these arborise mainly within the intermediate reticular nucleus (IRt). Scale bar = 500 μm.

**Figure S3**

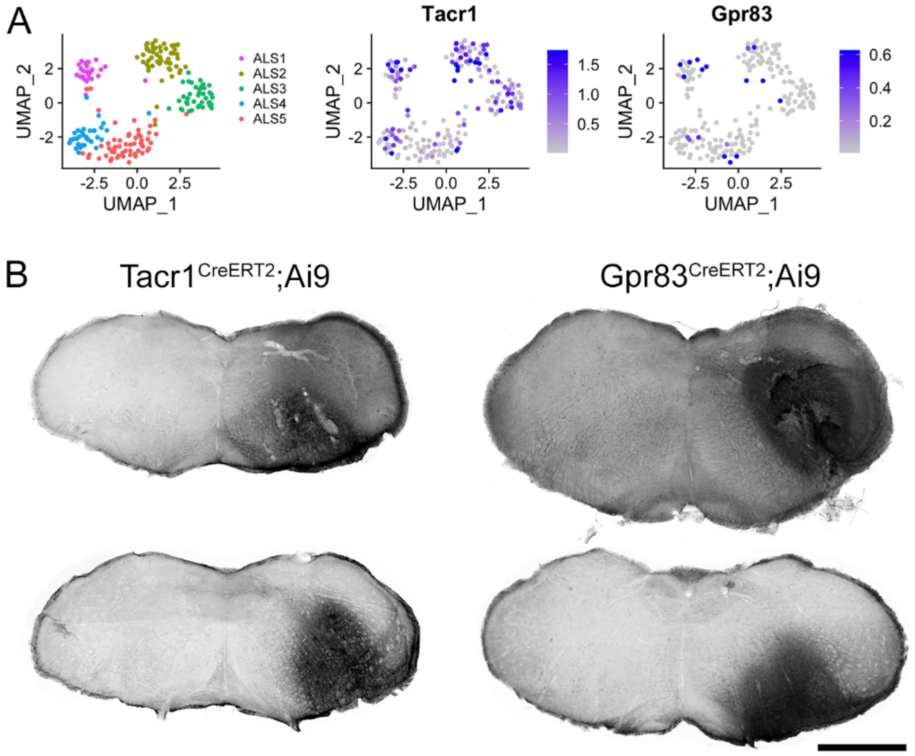


Expression of Tacr1 and Gpr83 by cells in ALS4. **A**: UMAP plot showing the distribution of cells in the ALS1-5 clusters, together with plots for expression of Tacr1 and Gpr83. Note the presence of many Tacr1-positive cells, and a few Gpr83-positive cells, in the ALS4 cluster. **B**: CTb injection sites in the four mice that were used to analyse expression of tdTomato in projection neurons in lateral lamina V. Scale bar = 1 mm.

**Figure S4**

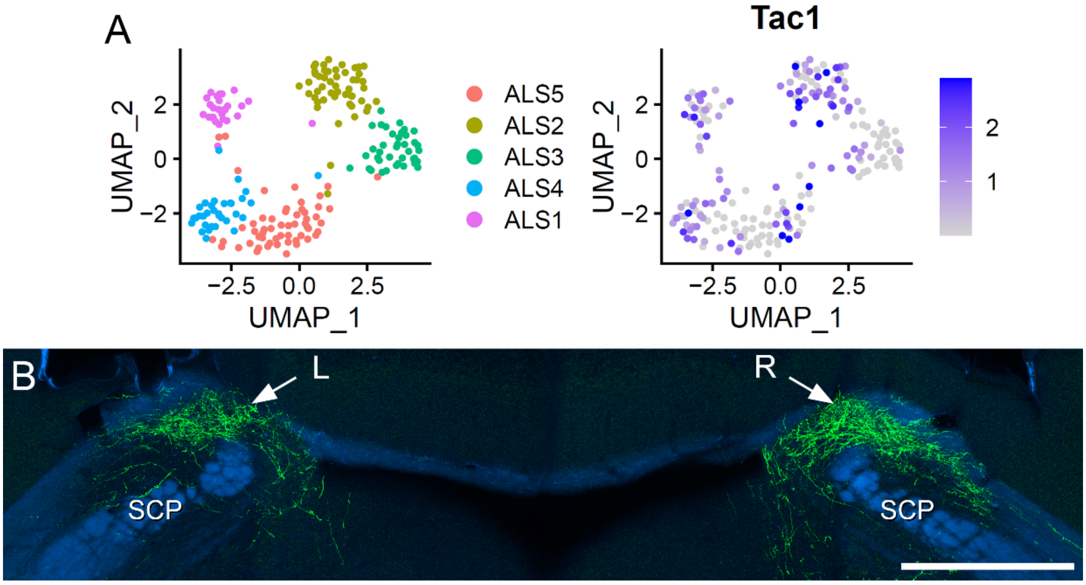


Expression of Tac1 by cells in ALS4. **A**: UMAP plot showing the distribution of cells in the ALS1-5 clusters, together with a plot for expression of Tac1. Note the presence of many Tac1-positive cells in the ALS4 cluster. **B**: anterograde labelling following injection of AAV.Cre$^{ON}$.GFP into the L5 segment on the right side of a Tac1$^{Cre}$ mouse. Immunolabelling for GFP (green) is superimposed on a dark-field image (blue). There is strong bilateral labelling with GFP (arrows) in the dorsalmost part of the lateral parabrachial area, corresponding to the PBil. SCP, superior cerebellar peduncle. Scale bar = 500 µm.

**Figure S5**

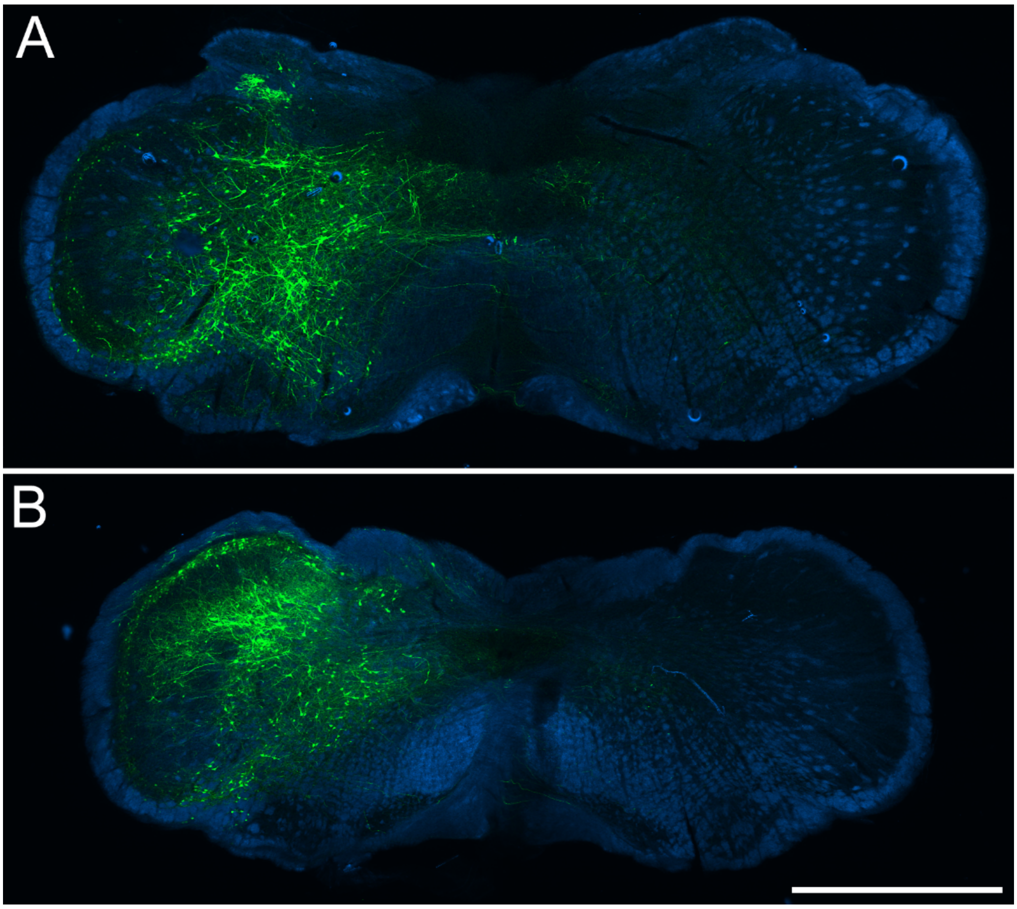


Injection sites for retrograde tracing of Sst-expressing ALS neurons. Sections through the medulla show the expression of GFP (green), superimposed on dark-field images (blue) following injection of AAV2r.Cre$^{ON}$.GFP into two Sst$^{Cre}$ mice. Note that GFP will only be expressed in Sst-expressing cells near the injection site. Scale bar = 1 mm.

**Table S1** Viruses used in this study

| | | | | | | Details of injection | |
|---|---|---|---|---|---|---|---|
| | **Construct(s)** | **Serotype** | **Promoter** | **Catalogue number** | **Source** | **Number of GCs** | **Volume** |
| AAV.Cre$^{ON}$.tdTom | tdTomato | AAV1 | CAG | v167-1 | VVF Zurich | $9.48 \times 10^7$ | 300 nl |
| AAV2r.Cre$^{ON}$.GFP | GFP | AAV2r | shortCAG | v158-retro | VVF Zurich | $2.15 \times 10^9$ | 500 nl |
| AAV.Flp | Flp | AAV9 | shortCAG | v453-9 | VVF Zurich | $1.24 \times 10^9$ | 400 nl |
| AAV.Cre$^{ON}$/Flp$^{ON}$.GFP | GFP | AAV1 | hEF1α | vHW33-1 | VVF Zurich | $1.62\text{-}8.1 \times 10^7$ | 300 nl |

The values in columns 7 and 8 refer to individual injections. GCs, gene copies; GFP, green fluorescent protein.

**Table S2** Antibodies used in this study

| Antibody | Species | Source | Catalogue # | Dilution | RRID |
|---|---|---|---|---|---|
| GFP | Chicken | Abcam | ab13970 | 1:1000 | RRID:AB_300798 |
| mCherry* | Rat | Invitrogen | M11217 | 1:1000 | RRID:AB_2536611 |
| CTb | Goat | List Biological | 703 | 1:5000 | RRID:AB_10013220 |
| NK1r | Rabbit | Sigma-Aldrich | S8305 | 1:1000 | RRID:AB_261562 |
| NeuN-Alexa647 | Rabbit | Abcam | ab190565 | 1:1000 | RRID:AB_2732785 |

*The mCherry antibody also recognises tdTomato
